# A connectome-based neuromarker of the non-verbal number acuity and arithmetic skills

**DOI:** 10.1101/2021.07.09.451740

**Authors:** Dai Zhang, Liqin Zhou, Anmin Yang, Shanshan Li, Chunqi Chang, Ke Zhou, Jia Liu

## Abstract

The approximate number system (ANS) is vital for survival and reproduction in animals and crucial in constructing abstract mathematical abilities in humans. Most previous neuroimaging studies focused on identifying discrete brain regions responsible for the ANS and characterizing their functions in numerosity perception. However, there lacks a neuromarker to characterize an individual’s ANS acuity, especially one based on the whole-brain functional connectivity (FC). Here, we identified a distributed brain network (i.e., numerosity network) using a connectome-based predictive modeling (CPM) analysis on the resting-state functional magnetic resonance imaging (rs-fMRI) data based on a large sample size. The summed strength of all FCs within the numerosity network could reliably predict individual differences of the ANS acuity in behavior. Furthermore, in an independent dataset from the Human Connectome Project (HCP), we found that the summed FC strength within the numerosity network could also predict individual differences in arithmetic skills. Our findings illustrate that the numerosity network we identified could be an applicable neuromarker of the non-verbal number acuity and might serve as the neural basis underlying the known link between the non-verbal number acuity and mathematical abilities.

## Introduction

The number of items in a set is represented in humans by two different representational systems: a language-dependent cultural system that encodes the precise cardinality of elements ^1^; and an approximate number system (ANS) shared by adults ^2, 3^, infants ^4^, many animal species ^5, 6, 7^, and even deep convolutional neural networks ^8, 9, 10^, which encodes quantity in an approximate, non-symbolic manner without verbally counting (i.e., numerosity perception). In animals, the ANS is of evolutionary importance as it can guide behaviors necessary for survival and reproduction, such as determining the relative amount of food during foraging ^11, 12^. In humans, the ANS is hypothesized to be the foundation of constructing more abstract mathematical abilities ^13^. For example, Harvey *et al*. ^14^ revealed a positive correlation between ANS acuity and math achievement, indicating the importance of ANS in shaping symbolic math skills.

Over decades, significant progress has been made to understand how numerosity information is represented in the brain. For instance, neurophysiological studies revealed that neurons in the prefrontal and parietal cortices of macaques were tuned to the numerosity of spatial arrays ^15, 16, 17, 18^. In humans, using the fMRI approach, researchers have also found that the numerosity information was mainly represented in the frontoparietal association cortex, including the horizontal part of the intraparietal sulcus, which exhibited similar tuning to numerosity ^19, 20, 21, 22^ (add He et al., PNAS,2015). However, recent new evidence showed that other brain areas were also involved in processing numerosity information. For example, Harvey ^23^ has proposed a numerical neural system including visual cortex, inferior temporal gyrus (ITG), fusiform gyrus, and angular gyrus, in addition to the parietal lobe and prefrontal cortex. The earliest numerosity-related neural activity has also been found in the visual cortex, such as V2 and V3 ^24, 25^, and numerosity information could be decoded in both the early visual and intra-parietal cortex ^26^. Number neurons were also found in the medial temporal lobe (MTL) of human neurosurgical patients when they performed calculation tasks on symbolic and non-symbolic stimuli ^27^. The angular gyrus also plays a central role in the linguistic number representation system ^28, 29^. Taken together, the neural representation of the ANS system seems to involve multiple cortical areas. However, most of these previous studies focused on identifying the function of specific brain regions supporting the ANS. There still lacks a connectome-wise neuromarker based on whole-brain functional connectivity (FC) that can characterize an individual’s ANS acuity.

FC measures the activity synchronization or interaction between fMRI bold oxygen level-dependent (BOLD) signals of two brain areas ^30, 31^. Whole-brain FC analysis based on network theory provides information on how distributed brain regions simultaneously exchange or integrate information to support specific cognitive functions ^32^. Recently, a connectome-based modeling (CPM) analysis ^33^ has been proposed to predict individual differences in human behavior (e.g., cognitive abilities) based on participants’ whole-brain FC pattern. The CPM analysis provides a complementary measure of human cognitive abilities to traditional approaches that focused on specific functions of discrete brain regions. The CPM approach can be successfully applied to predict individual differences in several cognitive abilities, such as fluid intelligence (gF) ^34^, sustained attention ^35^, reading accuracy ^36^, and creative ability ^37^. Because substantial variations also exist in the ANS acuity among individuals ^14, 38, 39^, here in the present study, we aimed to employ the CPM analysis to develop a whole-brain neuromarker (i.e., numerosity network) for the ANS acuity based on the resting-state fMRI data.

In addition, previous studies also illustrated that the CPM identified with one task could be generalized to predict the performance of other related tasks. For instance, sustained attention CPM, derived from healthy individuals in a sustained attention task, could also predict the ADHD-Rating-Scale (ADHD-RS) score in the ADHD patients ^35^, and the reading recall accuracy in another group of healthy individuals ^36^. As mentioned above, because the ANS plays a crucial role in determining an individual’s math achievement ^14^, the second aim of the present study is to examine whether the numerosity network that we identified above could also predict an individual’s arithmetic skills.

Specifically, in a large sample dataset (n>250), we measured each individual’s ANS acuity (i.e., numerosity precision) using a dot-array number comparison (NC) task ^14^. We then constructed a numerosity CPM based on their resting-state fMRI data to predict the individual differences in ANS acuity, using a leave-one-out (LOO) cross-validation procedure. As a result, the summed FC strength within the numerosity network could predict individual differences of the ANS in the left-out participant. More importantly, we further examined whether the numerosity network can predict the performance of arithmetic skills in an independent dataset from the Human Connectome Project (HCP). Our results suggested that the summed FC strength within the numerosity network could also predict individual differences in arithmetic skills, not language comprehension abilities. In sum, our findings demonstrate that the ANS acuity can be reliably predicted from the strength of the numerosity network of each individual, and imply that the numerosity network might also serve as the neural basis underlying the known link between ANS acuity and mathematical ability.

## Results

### Behavioral performance

The present investigation included an ANS dataset and an HCP math/story dataset. The ANS dataset consisted of a primary dataset and a validating dataset. It was used to obtain and verify the numerosity network. The HCP math/story dataset was used to investigate whether the numerosity network identified in the previous ANS dataset could predict individual differences in arithmetic skills and/or language comprehension abilities. In the ANS dataset, we calculated the Weber fraction for each participant, representing the participant’s ANS acuity ^14^. In the HCP math/story dataset, we calculated two measures of the inverse efficiency score (IES) for each participant, representing the participant’s arithmetic skills and language comprehension abilities, respectively ^40^. Larger scores of Weber fraction or IES performance indicate poorer numerosity acuity or arithmetic skills, respectively. The mean performance for these behavioral tasks of all datasets was shown in Table 1.

**Table 1.**
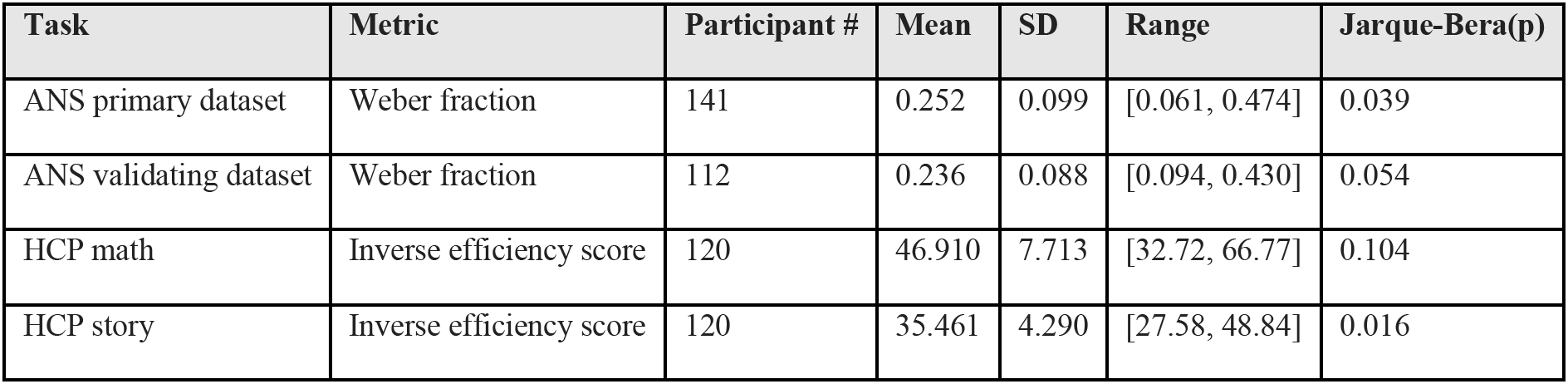
Summary of behavioral data. Jarque-Bera tests indicated significant departures from normality, supporting the use of Spearman’s rank correlations. Because we sought to identify any potential departures from normality, no correction for multiple comparisons was applied across these tests. All p-values are based on two-tailed tests.

Because the Jarque-Bera tests of normality revealed that these behavioral scores were not normally distributed (Fig. 1 B-E), we thus used Spearman’s rank correlation for the edge selection in establishing the numerosity network during the CPM analysis. Note that the numerosity network was constructed using the ANS primary dataset and validated using the ANS validating dataset.

**Fig 1.**
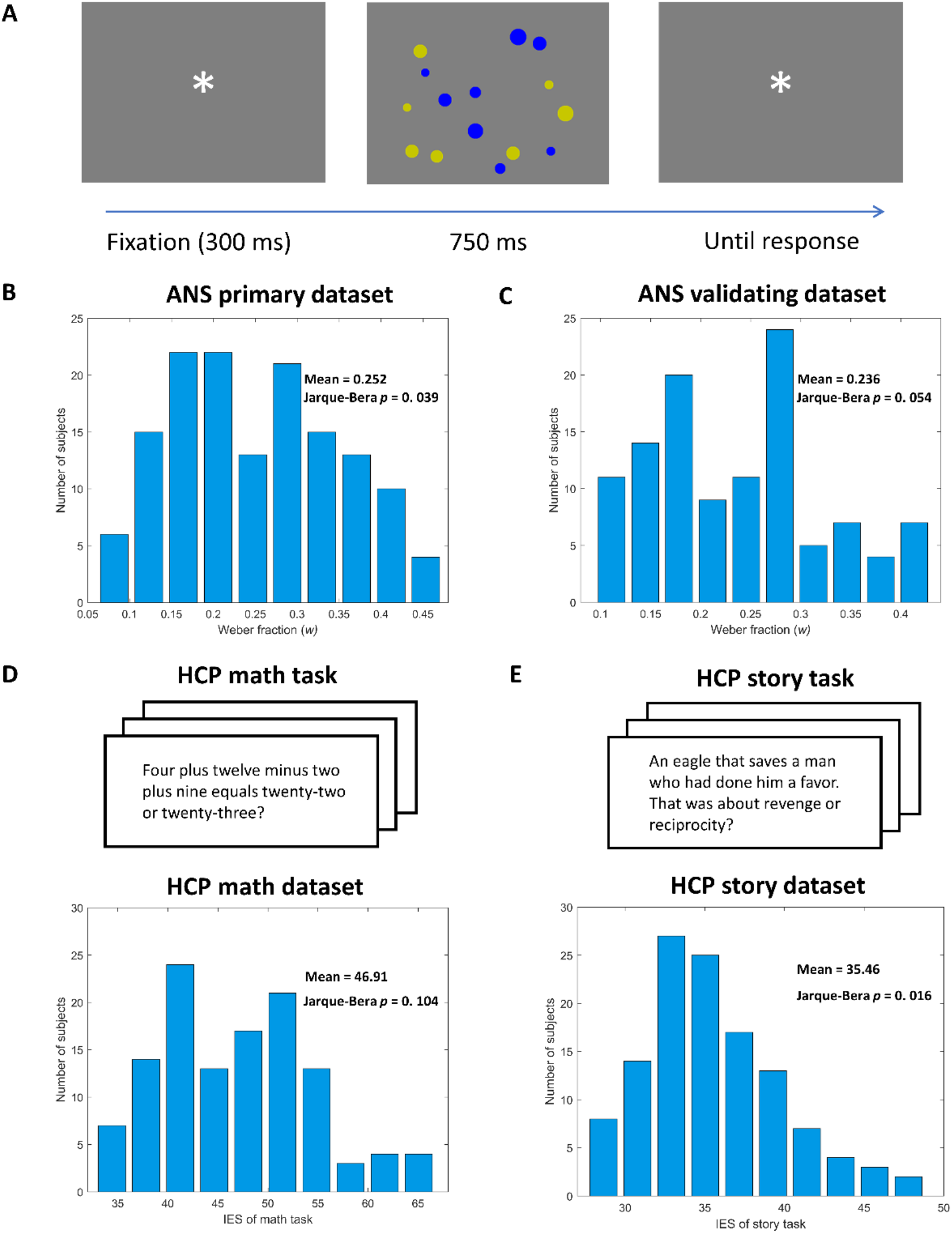
Experimental task and group performance for the ANS dataset and HCP story/math dataset. (A) The paradigm for the NC task. (B) Histogram of weber fraction w, the ANS acuity, for the ANS primary dataset (n=141), as determined by a psychophysical model for each participant. (C) Histogram of weber fraction w for the ANS validating dataset (n=112). (D/E) Experimental task and histogram of the IES performance for the HCP math/story dataset (n=120).

### Numerosity network can predict the individual differences in the ANS acuity

In the ANS primary dataset (141 participants), using the Shen *et al*. protocol ^33^, we adopted the LOO cross-validation method to test whether the intrinsic FC profile can predict the Weber fraction. For each LOO iteration, all 141 participants were divided into training (140 participants) and testing (1 left-out participant) sets. In the training set, for each edge between pair of nodes within the Shen’s atlas ^41^, we calculated the correlation between its FC and the Weber fraction across participants. Edges showing significant positive or negative correlation (*p*<0.01) composed a positive or negative predictive network, respectively. Next, for each of the 140 participants, we calculated the summed strength of all FCs within the positive and negative networks separately. We constructed a general linear model (GLM) for both the positive and negative networks by relating the summed strength to the Weber fraction across 140 participants in the training set. Then, in the testing set, the GLM was used to obtain the predicted Weber fraction of the left-out participant from his/her summed FC strength within the positive and negative network separately.

Across all 141 LOO iterations, we then calculated the Pearson’s correlation between the observed and predicted Weber fraction scores to evaluate the predictive power. For the positive networks, the correlation between observed and predicted Weber fractions was significant (*r* = 0.204, *p* = 0.015), while for the negative networks, the correlation was non-significant (*r* = 0.032, *p* = 0.703) (Fig. 2). We carried out a permutation test to further confirm the reliability of these results. We shuffled the behavior performance across participants 10,000 times and calculated the Pearson’s correlation coefficients between observed (randomly shuffled) and predicted Weber fraction. We found that the positive network outperformed the set of 10,000 permutation tests (*p*_perm_ = 0.0081). In comparison, the negative network did not outperform the set of 10,000 permutation tests (*p*_perm_ = 0.3510). There was a significant difference in the predictive power between the positive and negative networks (Steiger’s *z* = 2.67, *p* = 7.6×10^-3^) ^42^. It thus suggests that the strength of the functional connectivity profile within the positive networks could predict individual differences in ANS acuity. However, the negative network did not show the predictive ability of the individual differences in ANS acuity. Therefore, in the subsequent analysis, we focused on the predictive power of the positive network. Note that the predictive ability of the numerosity network cannot be alternatively explained by head motion, as the average frame-to-frame motion was not correlated with Weber fraction (*r* = −0.0149, p = 0.8607).

**Fig 2.**
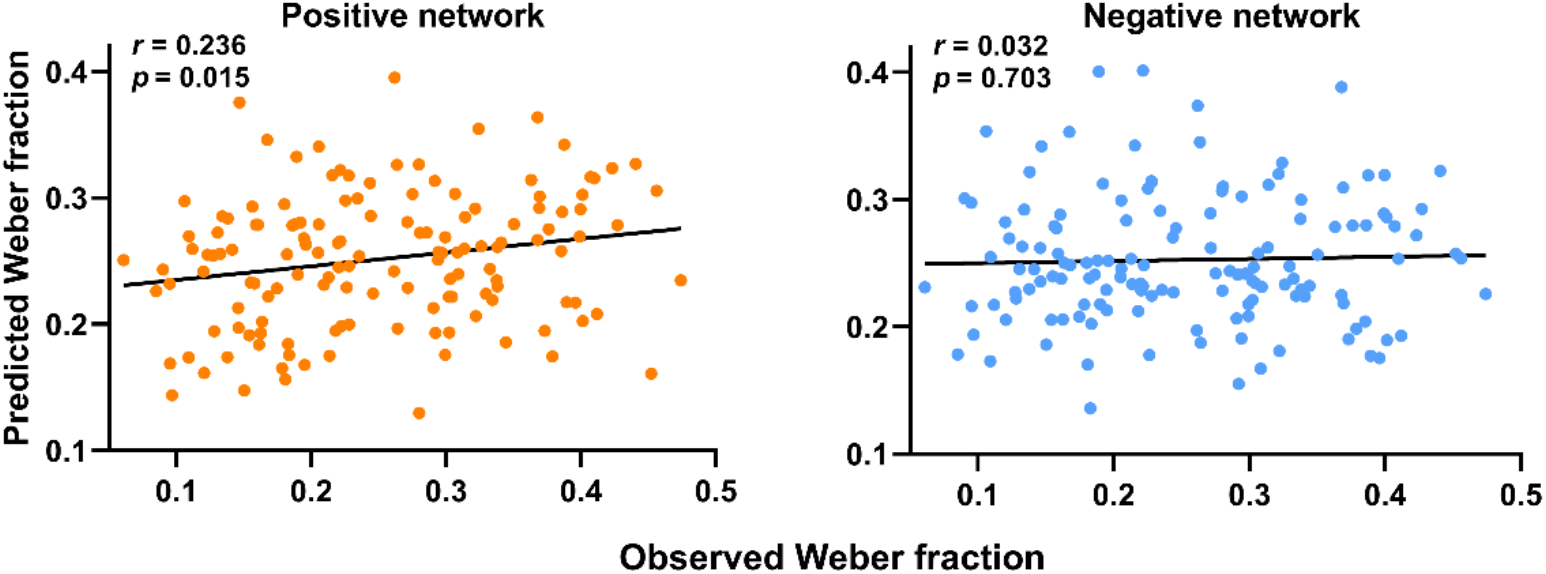
CPMs predict Weber fraction. Scatter plots show correlations between observed Weber fraction and predictions by positive (left) and negative (right) networks. Network models were iteratively trained on behavior and MRI data from n-1 participants in the ANS primary dataset and tested on the left-out participant. The *r* and *p*-value above each plot show the Pearson’s correlation coefficient between observed and predicted Weber fraction, and corresponding significance level.

Note that the positive networks differed across iterations. Across all 141 iterations, the number of edges within the positive ranged from 114 to 155. We generated a final numerosity network by selecting the overlapping edges of the positive networks across all iterations. There were 87 nodes and 80 edges in the final numerosity network.

Then, in the independent validating dataset (112 participants), we evaluated whether the final numerosity network could successfully predict the individual differences of the ANS acuity. The correlation between the summed strength of the numerosity network and the Weber fraction across participants was significant (*r* = 0.236, *p* = 0.012) (Fig. 3 A). We also conducted the same permutation test to confirm this result further. The numerosity network outperformed the set of 10,000 permutations where we randomly shuffled the behavior performance across participants (*p*_perm_ = 0.0060). It thus indicated that the final numerosity network we identified could serve as a connectome-based neuromarker to predict an individual’s ANS acuity reliably.

**Fig 3.**
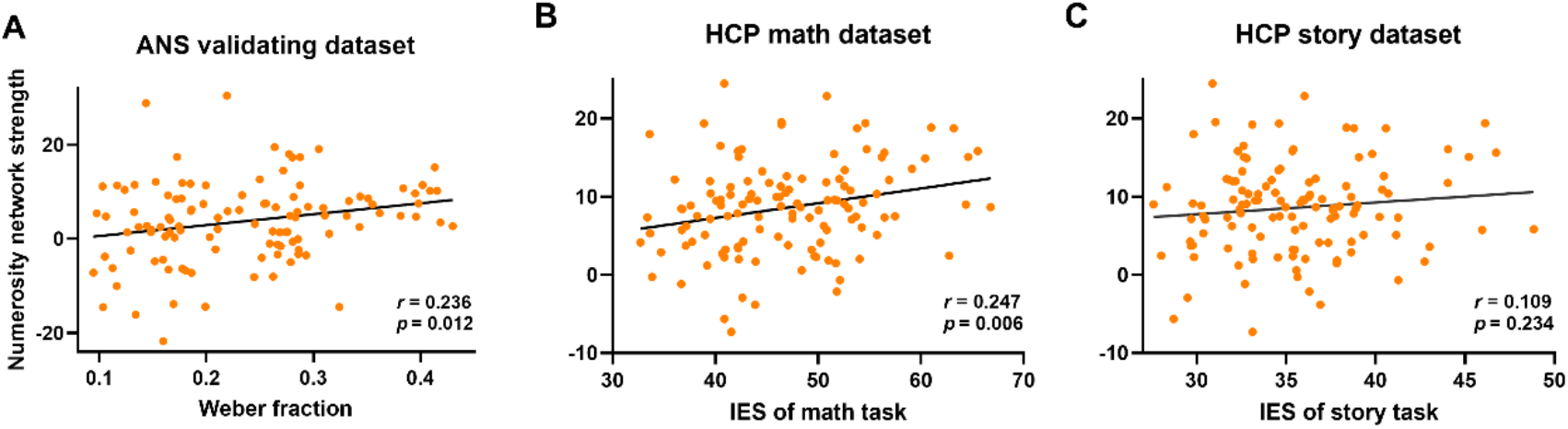
The predictive power of numerosity neuromarker on several independent datasets. Pearson’s correlations between network strength of numerosity network and behavior performance in the ANS validating dataset (A), HCP math dataset (B), and HCP story dataset (C) were calculated to assess the predictive power of numerosity neuromarker. The meaning of *r* and *p* shows the Pearson’s correlation coefficient between numerosity network strength and each behavior performance, and corresponding significance level, respectively.

### Summed FC strength within the numerosity network can predict arithmetic skills

To address our second aim, we further evaluated whether the summed FC strengths within the numerosity network could predict the individual differences in the arithmetic skills in an HCP math/story dataset. We found that the summed FC strengths within the numerosity network we identified above correlated significantly with the IESs in the math task (*r* = 0.247, *P* = 0.006) (Fig. 3 B). The numerosity network also outperformed the set of 10,000 permutation tests where we randomly shuffled the IES performance in the math task across participants (*P*_perm_=0.0025). The greater the summed FC strength within the numerosity network, the larger the IES measure of the arithmetic skills. Greater summed FC strength within the numerosity network pointed to both the higher Weber fraction (i.e., poorer numerosity precision) in the NC task and the larger IES (i.e., poorer arithmetic skills) in the math task. It thus indicates that the numerosity network could also predict the performance of arithmetic skills, even in a completely new dataset.

We next evaluated whether the numerosity network could predict the IES of the story task. We found that the summed FC strength within the numerosity network could not predict the IES in the story task (*r* = 0.109, *P* = 0.234) (Fig.3 C). We again ran permutation tests and found that the numerosity network failed to outperform the set of 10,000 permutations where we randomly shuffled the IES performance in the story task across participants (*P*_perm_=0.1161).

In short, these findings suggest that the numerosity network could only predict the individual differences in the arithmetic skills in specific, but not language comprehension abilities in general.

### Anatomical locations of network edges

We also examined the Anatomical locations of network edges within the numerosity network. The numerosity neuromarker involves a distributed neural network in the neocortex (Fig. 4B). As shown in Fig. 4A, in the numerosity network, 45 nodes were in the right hemisphere, and 42 nodes were in the left hemisphere. 18, 17, 13, 13, 12, 11, and 3 nodes located in seven macroscale regions (i.e., the temporal, prefrontal, motor, limbic, occipital, parietal, and insula) in Shen’s atlas ^41^ (Fig. 4C). Contralateral connections (50 edges) were more common than ipsilateral connections (30 edges) in the numerosity network (*χ*^2^(1) = 5.0022, *p* = 0.0253). There were 19 right-right connections and 11 left-left connections in the numerosity network (*χ*^2^(1) = 2.3793, *p* = 0.1229).

**Fig 4.**
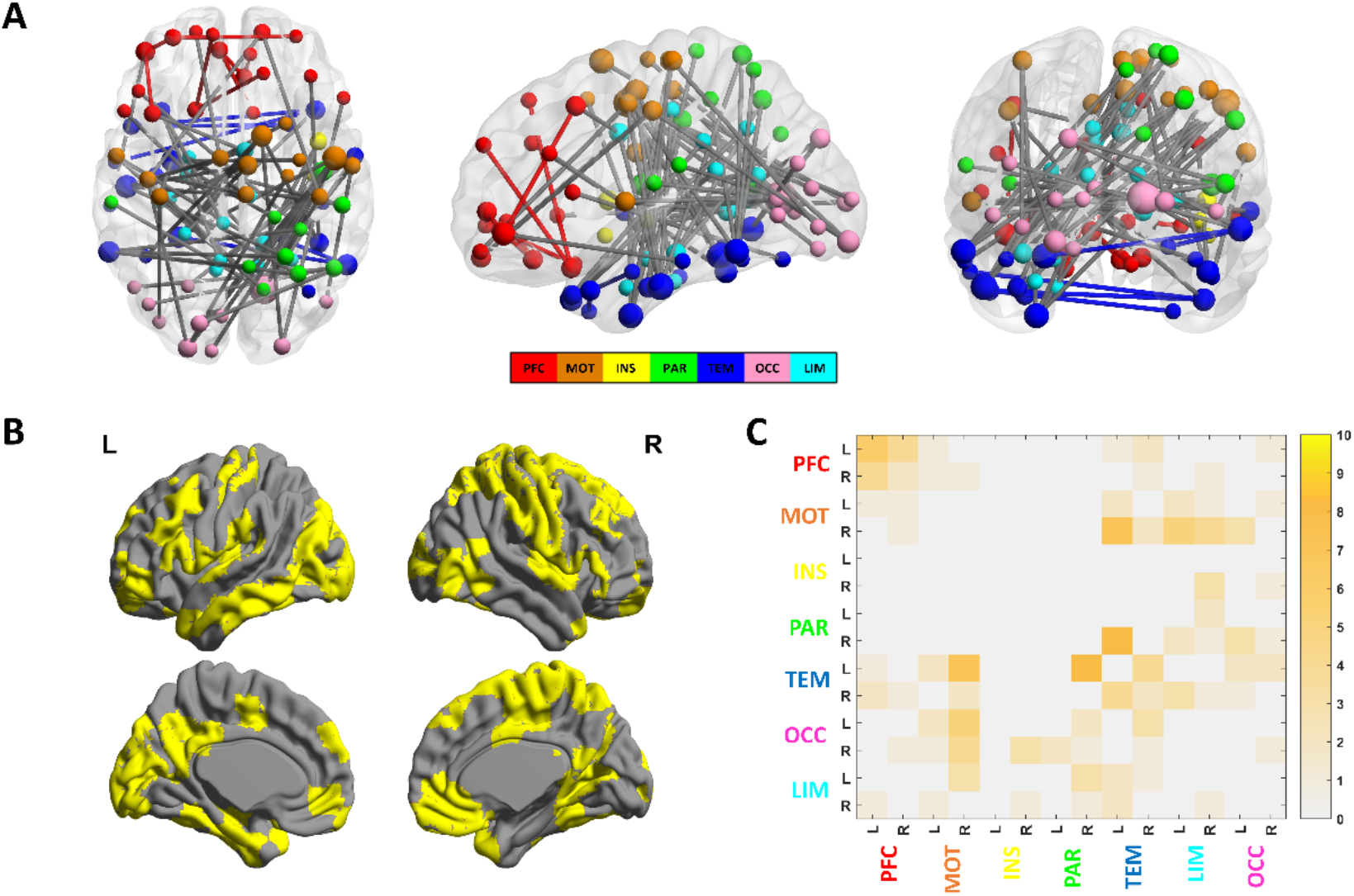
Anatomical locations of the numerosity network. (A) A 3d view of numerosity network edges and nodes on a glass brain. Nodes in the cerebellum, subcortical, and brainstem were not included in our analysis. Hence, Macroscale regions include the prefrontal cortex (PFC), the motor cortex (Mot), insula (Ins), parietal (Par), temporal (Tem), occipital (Occ), limbic (including the cingulate cortex, amygdala, and hippocampus; Lim). The larger size of a node indicates that the node gets a larger degree centrality (DC) value. (B) Cortical areas of the numerosity network. (C) All edges are grouped by macroscale region and hemisphere (L = left, R= right). Colorbar values indicate the number of edges within or between the macroscale regions.

The numerosity networks include many numerosity-related brain regions previously revealed, such as the early visual cortex ^43^, parietal sensory cortex ^26^, angular gyrus ^44^, medial temporal gyrus (MTG) ^27^, the motor cortex ^45^, and prefrontal cortex ^18^. However, some regions in the numerosity network, such as the visual association cortex, inferior temporal gyrus (ITG), and fusiform gyrus, have not been reported to be involved in processing numerosity information. The occipital (primary visual and visual association) nodes mostly connect to parietal and motor regions (Fig. 5A). There are dense connections between left temporal nodes and right motor, right parietal regions (Fig. 5B, 5C). The PFC nodes mainly connect within the lobes (Fig. 5D). Detailed information of all nodes and edges could be found in supplementary materials.

**Fig 5.**
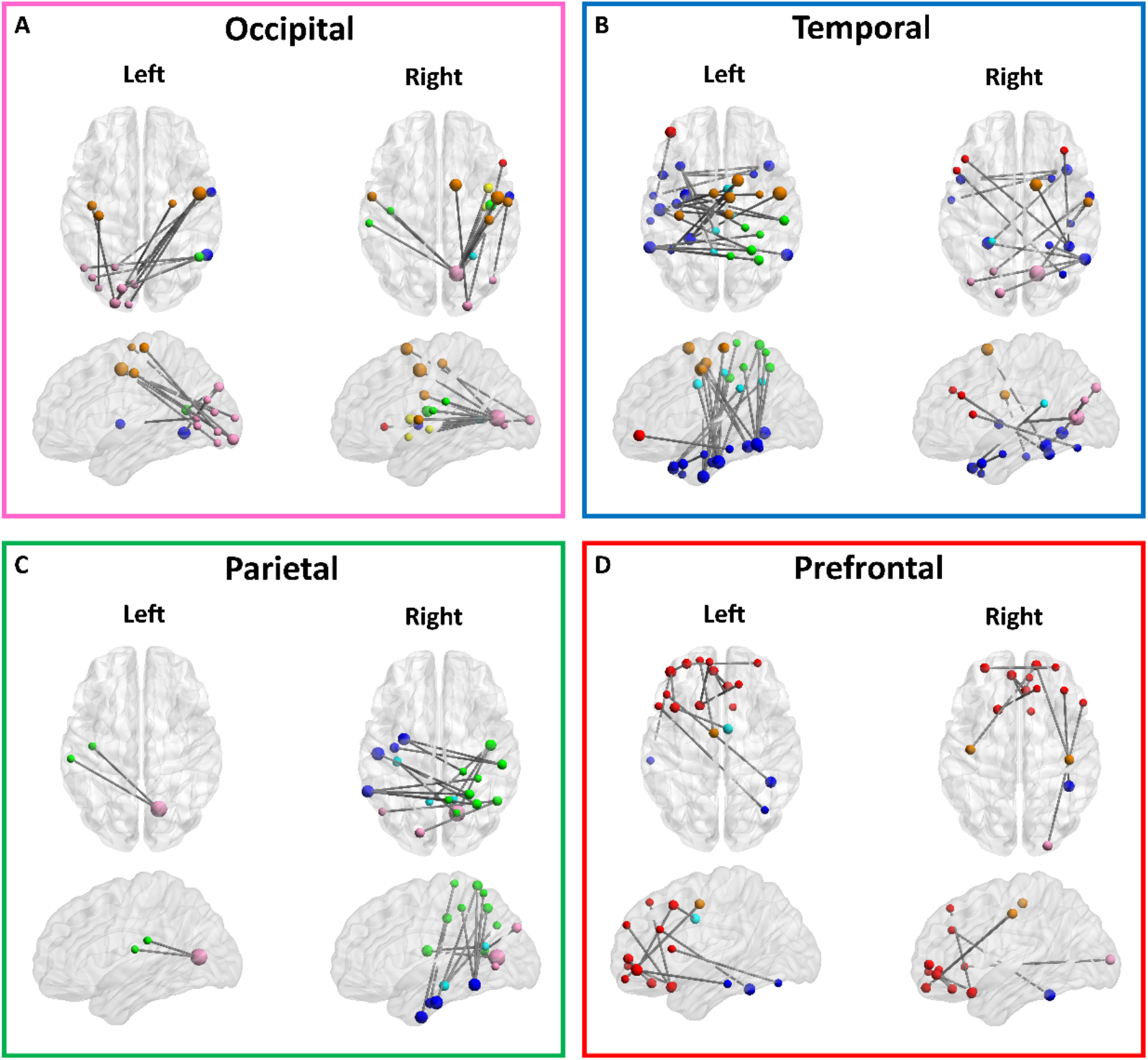
Functional connections in the numerosity network in each of four macroscale regions. (A) occipital, (B) temporal, (C) parietal, and (D) prefrontal regions. Dot color indicates macroscale region as in Fig. 4.

In addition, we investigated hub nodes in the numerosity network by ranking all nodes according to their degree centrality ^46^ (DC; i.e., the number of direct functional connections between a given node and the rest nodes within the numerosity network). The top 27 nodes with high DC are presented in Table 2 (DC>1). Regions with the highest DC indicates the hub regions of the numerosity network, including brain regions in the occipital (e.g., right primary visual cortex, Brodmann area (BA) 17, DC = 11), motor (e.g., right premotor cortex, BA 6, DC = 6), and temporal (e.g., left ITG, BA 20, DC = 5) cortex.

**Table 2.** Information of the hub nodes (DC>1) in the numerosity network.

| No. | DC | Node name | L/R | Macroscale region | BA | MNI |
| --- | --- | --- | --- | --- | --- | --- |
| 1 | 11 | Primary visual cortex | R | Occipital | 17 | (14.6, -68.3, 8.3) |
| 2 | 6 | Premotor cortex | R | Motor | 6 | (49.2, -4.6, 48.1) |
| 3 | 5 | Inferior temporal gyrus | L | Temporal | 20 | (-51.8, -18.2, -28.8) |
| 4 | 4 | Fusiform gyrus | L | Temporal | 37 | (-60.3, -50.0, -14.0) |
| 5 | 4 | Premotor cortex | R | Motor | 6 | (13.7, 6.3, 65.4) |
| 6 | 4 | Fusiform gyrus | L | Temporal | 37 | (-26.7, -42.7, -16.1) |
| 7 | 4 | Fusiform gyrus | R | Temporal | 37 | (55.2, -56.3, -4.8) |
| 8 | 4 | Parahippocampal cortex | L | Temporal | 36 | (-30.0, -5.8, -40.9) |
| 9 | 3 | Fusiform gyrus | R | Temporal | 37 | (41.6, -45.7, -22.6) |
| 10 | 3 | Frontopolar area | L | Prefrontal | 10 | (-42.7, 47.3, -6.9) |
| 11 | 3 | Temporopolar area | R | Temporal | 38 | (40.0, 18.9, -34.2) |
| 12 | 3 | Primary sensory cortex | R | Parietal | 1 | (43.3, -10.8, 13.9) |
| 13 | 3 | Supplementary motor area | R | Motor | 6 | (7.0, -8.1, 52.9) |
| 14 | 3 | Visual association cortex | L | Occipital | 18 | (-22.3, -96.6, -10.1) |
| 15 | 2 | Orbitofrontal cortex | L | Prefrontal | 11 | (-18.2, 19.0, -21) |
| 16 | 2 | Ventral anterior cingulate gyrus | R | Limbic | 24 | (5.3, -1.0, 35.6) |
| 17 | 2 | Angular gyrus | R | Parietal | 39 | (48.9, -58.1, 14.4) |
| 18 | 2 | Frontopolar area | L | Prefrontal | 10 | (-6.9, 48.3, -5.7) |
| 19 | 2 | Orbitofrontal area | R | Prefrontal | 11 | (5.1, 34.9, -17.4) |
| 20 | 2 | Primary sensory cortex | R | Motor | 1 | (42.0, -23.4, 53.4) |
| 21 | 2 | Paracentral lobule | R | Motor | 6 | (6.0, -22.3, 65.6) |
| 22 | 2 | Primary motor cortex | R | Motor | 4 | (57.8, -8.3, 27.3) |
| 23 | 2 | Inferior temporal gyrus | L | Temporal | 20 | (-49.3, -4.7, -37.4) |
| 24 | 2 | Visual association cortex | R | Occipital | 18 | (23.7, -96.0, 6.5) |
| 25 | 2 | Middle frontal gyrus | L | Prefrontal | 8 | (-39.3, 17.2, 46.7) |
| 26 | 2 | Inferior parietal gyrus | R | Parietal | 40 | (52.8, -27.2, 40.9) |
| 27 | 2 | Orbitofrontal cortex | R | Prefrontal | 11 | (13.9, 56.8, -16.6) |

BA, Brodmann area; DC, degree centrality; L, left; R, right.

## Discussion

In the present study, using the CPM analysis, we obtained a numerosity network based on the numerosity comparison task, whose strength could reliably predict the individual differences in the ANS acuity among individuals. This identified numerosity network consisted of functional connections between distributed brain regions, suggesting that a whole-brain, widely distributed numerosity network could serve as a reliable neuromarker for an individual’s non-symbolic numerosity ability. The robust predictive power of this numerosity neuromarker was further verified in a completely new dataset with the same behavioral task.

Only the positive network could predict individual differences in the ANS acuity. The negative network could not reliably predict the ANS acuity among individuals. Similar to our results, in predicting individual gF score ^34^, the negative network also showed less accurate predictions than the positive network. When the nodes were restricted within frontoparietal networks, the positive-feature models showed significant prediction results. In contrast, negative-feature models didn’t show significant predictive power ^34^. The reason for the inability of the negative network to predict the individual differences in the ANS acuity still needs future investigation.

Another significance of our study was that the numerosity neuromarker we identified could also specifically predict individual differences in arithmetic skills. In the literature of numerosity perception, many studies have found a positive correlation between numerosity precision and cognitive arithmetic skills ^14, 47^, suggesting that the number sense may serve as a ‘start-up’ tool for mathematics acquisition^13^. Our results provided a neural basis for this important link, suggesting that numerosity perception and arithmetic skills may share similar functional networks. On the contrary, this numerosity network could not predict language comprehension abilities, consistent with previous findings that mathematical processing relies on specific brain areas and dissociates from language processing ^48, 49, 50^.

The increased strength of the numerosity network was indicative of poor numerosity acuity or arithmetic skills. It seems to be counterintuitive. However, there were two possible explanations for this finding. One possible explanation was that this result might reflect a kind of compensatory mechanism ^51^. The participants with poor numerosity acuity need more sensory representation of the stimulus. Another possible explanation was that the strength of the numerosity network might represent the control ability to inhibit non-numerical magnitude information when extracting numerosity information, which can characterize numerosity information more accurately ^52, 53, 54, 55, 56^. However, our current findings cannot provide direct evidence to support any of these two accounts. Therefore, further investigations are needed to address this issue.

At the node level, the numerosity network involves multiple distributed regions, such as occipital regions (early visual cortex, visual association), sensorimotor regions (premotor cortex, supplementary motor cortex, primary sensory cortex), temporal regions (ITG, temporopolar area, and fusiform), and prefrontal areas (Frontopolar area and orbitofrontal cortex). Most of these regions have been suggested to involve in numerosity processing. For instance, Previous studies show that numerosity processing involves at least two temporal stages in the visual cortex ^57^. Subsequent studies found that the earliest activity likely arises from areas such as V2 and V3 ^24^. The sensorimotor cortex is suggested to be involved in a generalized system with quantitative information processing ^45^. In monkeys, neurons in the parietal sensorimotor area respond to the number of self-generated motor actions ^58^. A link between mathematics acquisition and motor skills has also been reported in children with learning disabilities, showing a correlation between motor skills and proficiency in solving mathematical problems ^59^. Both premotor and parietal areas get additional activation during verbal counting of visual and auditory items ^60^. Studies in humans ^61, 62^ indicated parts of the parietal cortex as a core number system that processes symbolic and non-symbolic numerical magnitude. There is also evidence that single-neuron activity from the medial temporal lobe (MTL) encoded numerical information when participants performed calculation tasks on symbolic and non-symbolic stimuli ^27^. Neurons in the lateral prefrontal cortex responded selectively to a specific number of items (i.e., numerosity) in visual multiple-dot displays ^63^. Therefore, the numerosity system might rely on the cooperative work of multiple distributed brain regions.

At the edge level, in the numerosity network, occipital regions are connected with angular gyrus, ITG, and fusiform gyrus, which could be conceptualized as a communication process between visual input and visual information extraction, such as visual recognition, spatial orientation, semantic representation processing. Moreover, the numerosity network did include many connections between occipital nodes and sensorimotor nodes, indicating the interaction between visual perception, action planning, and execution in the numerosity system. These results show a crucial role of the sensorimotor cortex in the numerosity system and the strong interaction between different perceptual features in the numerosity system ^45^. In addition, the numerosity network shows dense connections between left temporal and right premotor, and between left temporal and right parietal sensorimotor regions. These connections may present functional coupling between ventral and dorsal areas of the numerosity system. The numerosity network also shows connections within the prefrontal lobe. A nonverbal quantification system resides in a dedicated parietal-frontal brain network in primates ^64^. However, our numerosity network did not contain any parietal-frontal connections. Our findings provided a complementary measure of numerosity perception and arithmetic skills to traditional approaches that focused on the specific function of discrete brain regions.

However, it remains an open question whether the numerosity network we identified could predict changes in ANS acuity during development. Previous research has revealed that the precision of numerical acuity is sharped with age and the acquisition of formal mathematical education, which may reflect an ability to focus on numerical information while filtering out non-numerical information ^53, 54, 55, 56, 65^. Testing the change of the numerosity network associated with the within-individual changes in numerosity perception and mathematical abilities over years can inform the common and distinct functional architecture of these processes during development. In particular, it can provide insights into how functional brain organization reflects risk for or resilience to impairments such as developmental dyscalculia or mathematical difficulties, potentially informing early treatments or interventions.

A limitation of our study is that the ANS acuity and arithmetical skills were only assessed by specific paradigms. And we didn’t include the investigation of symbolic number representation. Further research could include various number-related tasks to verify our findings or address the common and distinct functional architecture of these tasks related to number perception and mathematical abilities.

## Methods

### Overview

In the primary ANS dataset, we constructed a numerosity CPM based on their behavior performance in an NC task and their resting-state fMRI data, to predict the individual differences in the ANS acuity, using a leave-one-out cross-validation procedure. The reliability of the identified numerosity network was verified in an independent validating ANS dataset with the same NC task. Then in the HCP math/story dataset, we examined whether the identified numerosity network could predict the individual differences in the arithmetic skills.

### ANS dataset

#### Participants

The behavioral data, and the structural and resting-state fMRI data were collected from two batches of college students (Batch A and Batch B). Data from the Batch A (154 participants; age: 20–25; mean = 21.66, SD = 1.07; 65 males) was used to construct the numerosity CPM. Data from the Batch B (145 participants; age: 20–25, mean = 21.39, SD = 0.91; 36 males) was used to validate the reliability of the numerosity network. Batch A and Batch B datasets correspond to the ANS primary and validating datasets, respectively. All participants have no history of neurological disorder (e.g., mental retardation, traumatic brain injury) or psychiatric illness. The experimental protocol was approved by the Institutional Review Board of Beijing Normal University. Written informed consent was obtained from all participants before the study.

#### NC task

Similar to previous research ^14^, a classical NC paradigm was used to measure participants’ ANS acuity. In each trial, a spatially intermixed blue and yellow dot display was presented on a computer screen for 750 ms (Fig1. A). Participants indicated which color was more numerous by the keypress. The ratio between the two sets of colored dots varied at 11 levels, including 12:11, 11:10, 10:9, 9:8, 8:7, 7:6, 6:5, 10:8, 8:6, 9:6, and 12:6. The color of the dots set was counterbalanced across trials. Dot-size of each ratio level was controlled: the average blue dot size variation was equal to the variation range of the average yellow dot size. Participants completed forty test trials after practicing five practice trials.

The minimum distinguishable difference between numbers of blue and yellow dots that produces a noticeable response was estimated as each participant’s ANS acuity known as the Weber fraction. The Weber fraction of each participant was estimated by a QUEST routine ^66, 67, 68^. The QUEST routine provides a given number of sequential trials and updating the probability distribution function (PDF) of Weber fraction based on the participant’s response and current PDF, following the Bayes’ Rule. After the final trial, the mean value of PDF was recorded as the participant’s Weber fraction and used to represent the participant’s ANS acuity. Thus, a greater Weber fraction score corresponds to the poorer ANS acuity.

Participants with divergent estimating sequences were excluded by visual inspection. Participants with Weber fraction greater than 0.5 were also excluded from the subsequent analysis (e.g., Weber fraction greater than 0.5 means an inability to distinguish 12 dots from 6 dots). In sum, 13 participants in the ANS primary dataset and 25 participants in the ANS validating dataset were excluded from further analysis.

#### MRI data and preprocessing

The structural and resting-state fMRI data were acquired in the same session. The MRI data were acquired from a Siemens 3T Trio scanner (MAGENTOM Trio, a Tim system) with a 12-channel phased-array head coil at the BNU Imaging Center for Brain Research, Beijing, China. T1-weighted structure images were acquired with a magnetization-prepared rapid gradient-echo (MPRAGE) sequence (TR/TE/TI = 2.53 s/3.45 ms/1.1 sec, FA = 7 degrees, voxel size = 1×1×1 mm, slice thickness = 1.33 mm, number of volumes = 128) for each participant. The resting-state data was acquired using a T2*-weighted GRE-EPI sequence with different parameters from task-state fMRI (TR = 2000 ms, TE = 30 ms, flip angle = 90 degrees, number of slices = 33, voxel size = 3.125 × 3.125 × 3.6 mm).

Resting-state fMRI images were preprocessed using the FMRI Expert Analysis Tool version 5.98, part of FMRIB’s Software Library (www.fmrib.ox.ac.uk/fsl). The first four volumes of each participant were discarded to allow for the stabilization of magnetization. In addition to head-motion correction, brain extraction, spatial smoothing (FWHM = 6 mm), grand-mean intensity normalization, and removing a linear trend, several other preprocessing steps were also used to reduce spurious variance. These steps included using a temporal band-pass filter (0.01–0.08 Hz) to retain only low-frequency signals, and regression of the time course obtained from motion correction parameters, the mean signals of the cerebrospinal fluid and white matter, and the first derivatives of these signals. Then, all functional images were aligned to the structural images using FMRIB’s linear image registration tools and warped to the MNI152 template using FMRIB’s nonlinear image registration tool. Since head motion might confound functional connectivity analyses, in the validating dataset, we excluded eight participants with frame-to-frame head motion estimation greater than 0.15 ^34^. Finally, 141 participants were retained in the ANS primary dataset, and 112 participants were retained in the ANS validating dataset. There was no correlation between head motion and Weber fraction in the retained participants of both datasets (the primary dataset: *r* = −0.0149, *p* = 0.8607; the validating dataset: *r* = −0.0068, *p* = 0.9433).

#### Functional connectivity calculation

FC was calculated between ROIs or “nodes” in Shen’s atlas, which maximized the similarity of the time series of the voxels within each node ^41, 69^. A subset of nodes of the 268-node functional brain atlas was used in our study because some resting-fMRI scans did not cover the full brainstem and cerebellum. We thus focused on nodes in the neocortex by removing all nodes of the cerebellum, brainstem, and subcortex (67 nodes totally). The remained 201 nodes were shown in Fig. S1. The mean time course of each node was extracted as a measure of spontaneous neural activity in that node. Pearson’s correlation coefficients (*r*) were calculated between the time courses of each pair of nodes and normalized using Fisher’s *r*-to-Z transformation. Finally, a 201×201 FC matrix was obtained for each participant in the primary dataset and the validating dataset.

### HCP dataset

#### Participants

Data for evaluating the numerosity network’s predictive power in predicting the individual differences in mathematical and language comprehension abilities were from the Human Connectome Project (Q1 and Q2 HCP data releases; 131 participants; age: 22–35; 30 males). Resting-state fMRI data of both left-right (LR) and right-left (RL) phase-encoding runs (HCP filenames: rfMRI_REST1) were included in subsequent analysis. Consistent with the preprocessing procedure of the ANS dataset, we also excluded seven participants with frame-to-frame head motion estimation greater than 0.15 (violated in either of LR or RL phase-encoding runs, HCP: Movement_RelativeRMS_mean).

#### Behavioral Test

In the HCP language math/story task, participants answered math-related and story-related questions after hearing auditory blocks. The HCP language task consists of two runs that each interleaves four blocks of math task and four blocks of story task ^70^. The math task engages participants’ attention continuously with mental arithmetic. The math task includes auditory trials and requires participants to complete addition and subtraction problems, followed by a 2-alternative forced-choice task. For example, “Four plus twelve minus two plus nine equals *twenty-two* or *twenty-three*?” The math task is adaptive to maintain a similar level of difficulty across participants. The story task includes brief and engaging auditory stories (5-9 sentences) adapted from Aesop’s fables. For example, after a story about an eagle that saves a man who had done him a favor, participants were asked the question of “That was about *revenge* or *reciprocity*?” For both tasks, participants push a button to select either the first or the second choice. Median reaction time and accuracy were recorded for both math and story tasks (HCP math task: Language_Task_Math_Acc and Language_Task_Math_Median_RT, HCP story task: Language_Task_Story_Acc and Language_Task_Story_Median_RT).

We calculated the inverse efficiency score (IES) to integrate response time and accuracy ^40^. The IES was calculated as median response time divided by accuracy, to evaluate mathematical and language comprehension abilities. There was no correlation between head motion and IES in the remaining set of n = 124 participants for math and story tasks (LR: IES of Math r= 0.05, p= 0.5588, IES of Story r= −0.03, p= 0.76; RL: IES of Math r= 0.01, p= 0.9114, IES of Story r= −0.09, p= 0.32).

#### MRI data and preprocessing

The HCP minimal preprocessing pipeline was used for the HCP data set ^71^. This pipeline includes artifact removal, motion correction, and registration to standard space. After this pipeline, several standard preprocessing procedures, including linear detrend, temporal filtering (0.01-0.1Hz), regression of 12 motion parameters (HCP data; these include first derivatives, given as Movement_Regressors_dt.txt) and mean time courses of the white matter and CSF as well as the global signal, were applied to the fMRI data ^34^. No spatial smoothing was used in HCP MRI dataset preprocessing.

#### Functional connectivity calculation

All steps to construct resting-state FC networks were identical to those used in the ANS dataset. Data from both LR and RL phase-encoding runs were used to calculate connectivity matrices, respectively. The average of these two connectivity matrices was used as the participants’ connectivity matrices. Then a 201×201 FC matrix was obtained for each participant.

### Numerosity CPM and numerosity network

#### Numerosity CPM

Numerosity CPM was established to predict individual differences in the ANS acuity, using a leave-one-out (LOO) procedure ^35, 36^. One participant was left out for each LOO iteration as the testing set, and the remaining participants were regarded as the training set. The Spearman’s rank correlation was calculated across participants between functional connectivity of each edge in the FC matrix and Weber fraction. Consistent with previous research ^35^, for each LOO iteration, in the training set, the edges showing significant positive or negative correlation with a *p*-value below a threshold (*p*<0.01) across participants were included in a positive or negative network, respectively. The network strength of the training participant was defined as the participant’s summed strength of all FCs within the positive or negative network. A general linear model (GLM) was fit to relate the summed strengths to the Weber fraction for positive and negative networks, respectively. Then, in the testing set, the GLM was used to obtain the predicted Weber fraction of the left-out participant from his/her summed FC strength within the positive and negative network. After the LOO procedure was repeated for each participant iteratively, we calculated the Pearson’s correlation coefficient between the observed and predicted Weber fractions across all participants to evaluate the predictive power. The standard deviation (SD) method was applied to the Weber fraction to remove participants with an outlier of the Weber fraction. The Weber fraction of each participant was within the range of ± three times SD around the mean value. No participant was removed from the ANS primary dataset.

#### Permutation test

To confirm the reliability of prediction results, we fixed the predicted Weber fraction and shuffled the observed Weber fraction across participants 10,000 times. We calculated the Pearson’s correlation between predicted and observed (randomly shuffled) scores. The number of times that the correlation coefficients in the set of 10,000 permutation tests outperformed the correlation coefficient from the numerosity CPM divided by 10,000 was recorded as permutation test probability.

#### Numerosity network

Note that the positive or negative networks differed across iterations. To obtain a unique numerosity network, we generated a final numerosity network by selecting the overlapping edges across all LOO iterations’ network of the numerosity CPM (i.e., the overall network consisted of edges found in all LOO iterations). Then, the numerosity neuromarker was defined as the summed FC strength within the final numerosity network in this study. We evaluated the predictive power of the numerosity neuromarker on several datasets, including an independent ANS dataset and an HCP math/story dataset. Note that we used the numerosity network to denote the final numerosity network.

### The predictive power of numerosity network

#### The predictive power of numerosity network on ANS validating dataset

The ANS validating dataset was used to confirm the reliable predictive power of the numerosity network to an independent dataset with the same NC task. To evaluate the predictive power of the numerosity network, we calculated the Pearson’s correlation coefficients between the summed strength of the numerosity network and the Weber fraction of the ANS validating dataset. The standard deviation (SD) method (beyond the range of ± three times SD around the mean value) was used to exclude participants with outlier behavior performance from the correlation analysis. No participant was excluded from the ANS validating dataset.

#### Permutation test

To confirm the significance of results on the ANS validating dataset, we fixed the strength of the numerosity network and shuffled the observed Weber fraction 10,000 times, and calculated the Pearson’s correlation coefficients between them. Permutation test probability was calculated by the number of times that correlation coefficients in a set of 10,000 permutation tests outperformed the predictive power of the numerosity network, then divided by 10,000.

#### The predictive power of numerosity network on HCP dataset

The HCP (math/story) dataset was used to evaluate the predictive ability of the numerosity network to arithmetic skills and language comprehension abilities. The predictive power of the numerosity network is evaluated by Pearson’s correlation coefficients between the strength of the numerosity network and the behavioral performance of the HCP math/story dataset. Two participants (participant ID: 172332, 217429) in the HCP math task and two participants (participant ID: 193239, 255639) in the HCP story task were excluded because their performances were beyond the range of ± three times SD around the mean value. These four participants were excluded from the correlation analysis. Permutation test procedures were the same as those for the ANS validating dataset.

#### Anatomical distribution of numerosity network

To determine the anatomical location of the numerosity network, we grouped the 201 nodes into seven macroscale regions, including the prefrontal (PFC), motor (Mot), insular (Ins), parietal (Par), temporal (Tem), occipital (Occ), and limbic systems (Lim). We calculated the number of functional connections within or between all the seven macroscale regions. To investigate the hub nodes in the numerosity network, we ranked all nodes according to their degree centrality (DC; i.e., the number of direct functional connections between a given node and the rest nodes within the numerosity network).. The nodes and edges of the numerosity network were presented using BrainNet Viewer (V1.70, www.nitrc.org/projects/bnv) ^72^.

## Acknowledgments

This work was supported by the National Key R&D Program of China (2019YFA0709503), National Basic Research Program of China (2018YFC0810602), National Nature Science Foundation of China grant (31671133, 61971289), Shenzhen Science and Technology Research Funding Program (JCYJ20170412164413575, JCYJ20170412111316339), Fundamental Research Funds for the Central Universities, and Shenzhen-Hong Kong Institute of Brain Science-Shenzhen Fundamental Research Institutions.

## Data availability

Each participant’s functional matrix and behavior performance from the ANS dataset and HCP math/story dataset are available from the authors upon request.

## Code availability

Matlab scripts were written to identify numerosity neuromarker, make predictions from novel individuals’ FC matrices, and generalize the numerosity neuromarker to ANS validating dataset and HCP math/story dataset. These codes are available online from https://github.com/Dzhang1989z/Numerosity-CPM.

## Supplementary information

### Fine spatial distribution of numerosity network

The ROIs to which the occipital region is connected in numerosity network are visualized in Fig. 5A. The occipital (primary visual and visual association) nodes are mainly connected to temporal, parietal and motor regions. To be specific, the left occipital regions connect to right premotor cortex, left primary motor cortex, left primary sensory cortex, right angular gyrus, bilateral primary sensory cortex, right superior temporal cortex and right fusiform gyrus; the right occipital regions connect to right primary motor cortex, bilateral premotor cortex, bilateral primary sensory cortex, left supramarginal gyrus (SMG) and right superior temporal cortex.

The temporal regions played an outsize contribution in the numerosity network (Fig. 5B). The left temporal and right temporal region show sort of dissimilar connection patterns. The connection between left temporal (temporopolar area, medial temporal gyrus, inferior temporal gyrus and fusiform gyrus) nodes and right motor, right parietal regions is especially dense, while no connections between right temporal and parietal cortex was found. To be specific, the left temporal nodes mainly connected to several pre-motor nodes, right parietal sensory areas (located near the intraparietal sulcus, IPS) and bilateral ventral/dorsal posterior cingulate cortex (PCC); the right temporal nodes connected to a part of bilateral prefrontal cortex (PFC), right motor cortex, left temporal cortex and bilateral occipital cortex.

The ROIs connected to the parietal region are visualized in Fig. 5C. In the parietal region, most connections locate between right parietal sensory nodes and left temporal cortex, bilateral occipital cortex, as mentioned previously. The left parietal regions (primary sensory and SMG) merely connect with the primary visual area. The right parietal regions connect with the left inferior temporal gyrus (ITG), left fusiform gyrus, left parahippocampal gyrus (PPA), right primary visual cortex and left visual association cortex.

The right motor regions got more dense connections than the left motor regions (supplementary Fig. 2A). The motor (primary motor/sensory and premotor) nodes are mainly connected with occipital and temporal cortex. To be specific, the left motor regions connect with left ITG, left PPA, right primary visual and left visual association nodes; the right motor regions primarily connect with bilateral ITG, bilateral fusiform gyrus, left PPA, bilateral primary visual and visual association nodes.

The prefrontal cortex shows most connections located within prefrontal lobe (Fig. 5D). Besides the intra-lobe connections, the left prefrontal regions primarily connect with left premotor cortex, right fusiform gyrus and left medial temporal gyrus (MTG); the right prefrontal region connects to right primary sensory/motor cortex, right fusiform gyrus, right visual association area.

The limbic cortex shows connections with all other six macroscale regions excepting insula (supplementary Fig. 2B). Limbic cortex connects more densely with motor and parietal cortex than other cortices.

**Supplementary Fig. 1.**
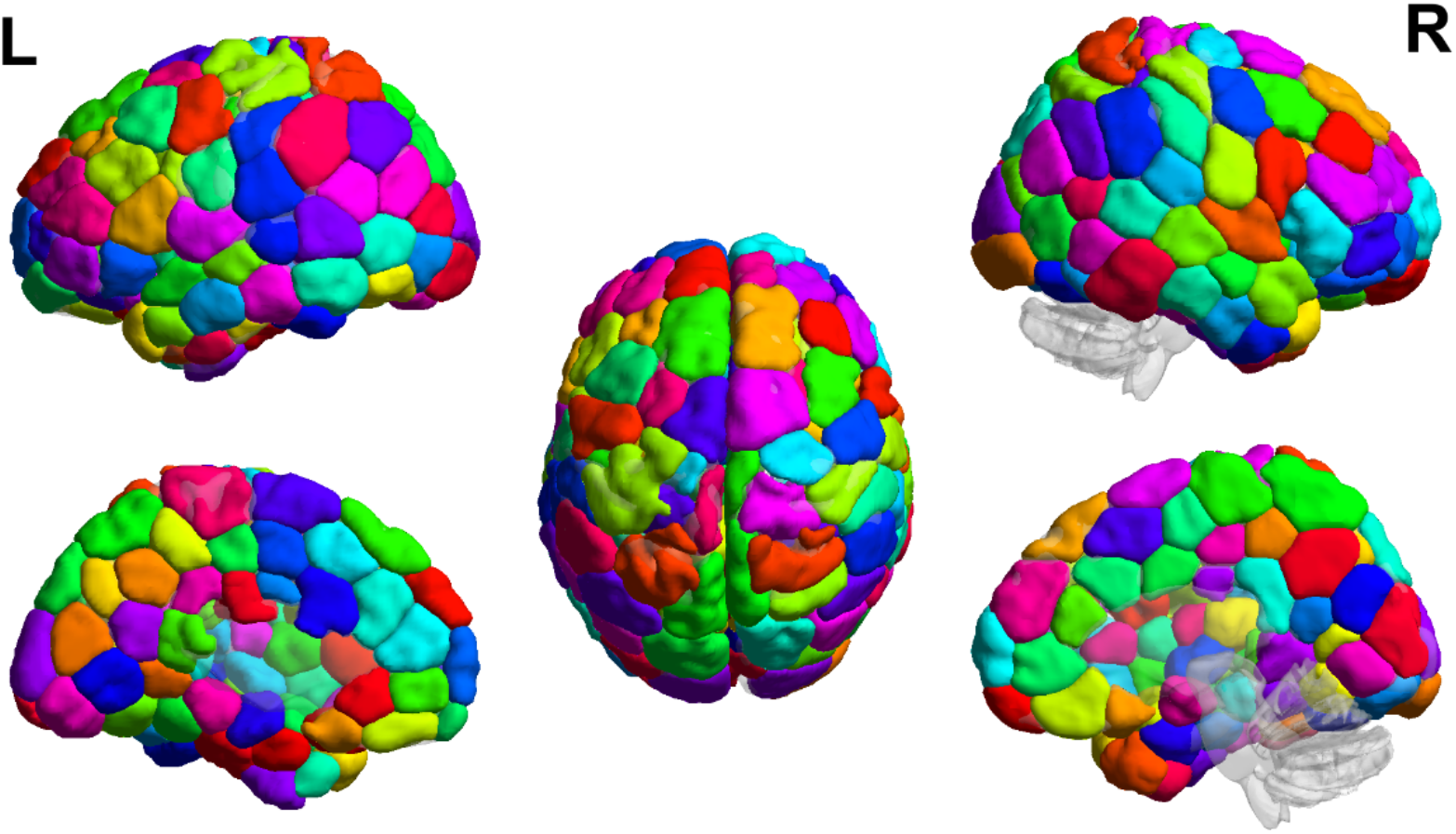
The 201-region functional parcellation used to define network nodes in primary NC dataset. Colored nodes were included in the FC matrix calculation. All nodes (67 nodes) in the cerebellum, brainstem, and subcortex were not included in our analysis.

**Supplementary Fig. 2.**
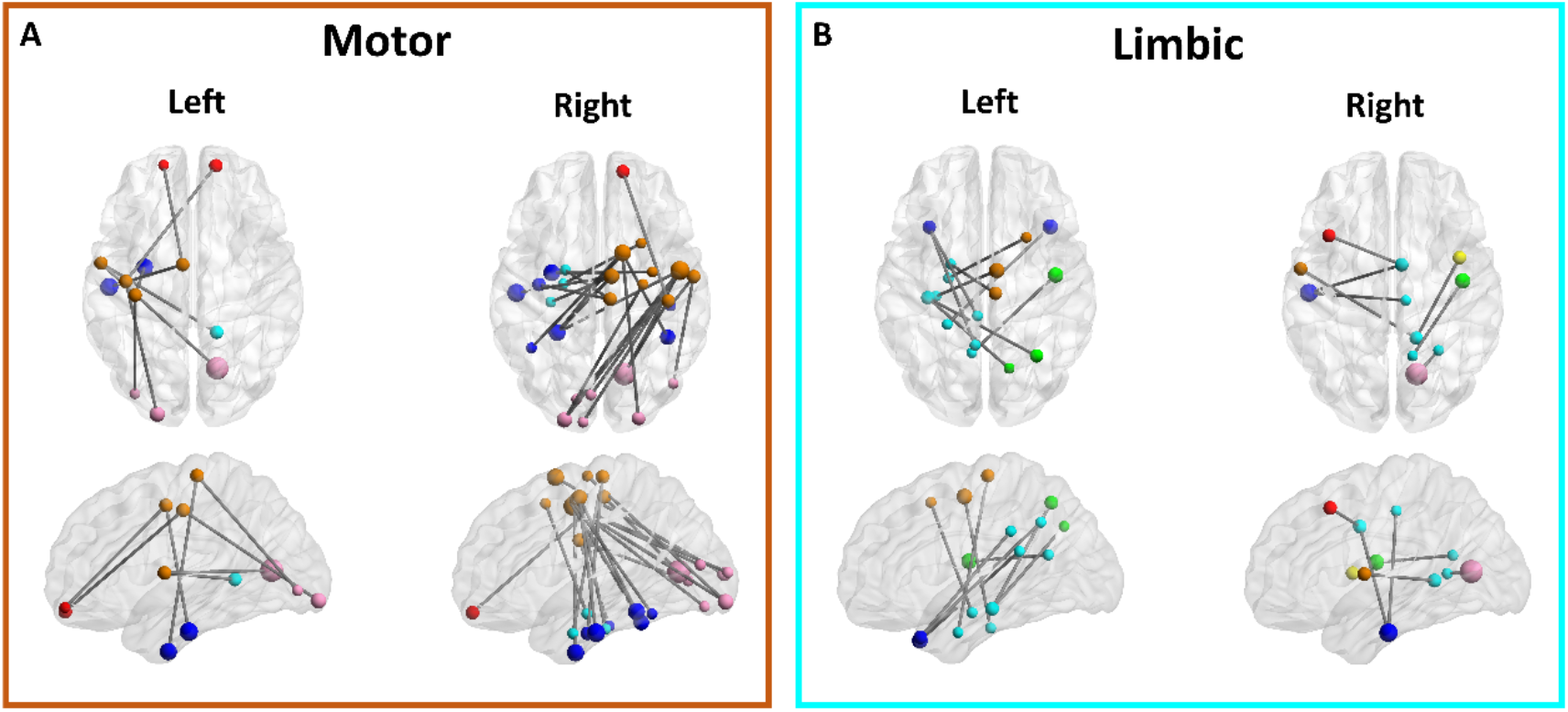
All connections in the numerosity network that include nodes in (A) motor and (B) limbic system. Dot color indicates macroscale region as in Fig. 4.

